# Active Learning framework for Antibody Binding Affinity Prediction

**DOI:** 10.64898/2026.10.17.759787

**Authors:** Chundru Sharath Krishna, Harshit Singh, Satya Pratik Srivastava, Rajeev Kumar Singh

## Abstract

Antibody engineering campaigns must prioritise variants from large combinatorial libraries under limited assay budgets. We present a Gaussian process (GP) active-learning benchmark for antibody-antigen binding affinity prediction across eight public AbBiBench assays spanning influenza hemagglutinin antibodies CR6261 and CR9114 and AAYL-series scFv libraries against the SARS-CoV-2 Spike HR2 peptide. The framework embeds variable heavy and light (VH/VL) sequences with ESM-2 650M, trains GP surrogates over compressed protein-language-model representations, and evaluates kernel, acquisition, representation, classical featurisation, and labelling-budget choices under a unified budget of 10% of each full pool with a label-blind seed and a fixed held-out evaluation split. Across the benchmark, Tanimoto+UCB has the highest mean Recall@2% point estimate among tested kernel-acquisition configurations, achieving **57.8%** mean Recall@2% across five random seeds (68.3% when scored against the acquisition-eligible pool), compared with 9.6% for random selection – a gap that is significant by a paired Wilcoxon test across datasets (*p* = 0.0078); its point estimate is not significantly different from RQ+UCB or Tanimoto+EI at five seeds. We adopt the continuous Tanimoto kernel of Gessner et al. (2024). The same configuration recovers the largest fractions of top binders in the influenza libraries (74.2% on `3gbn_h1`, 82.7% on `3gbn_h9`, and 84.9% on `4fqi_h3`) while exposing harder AAYL combinatorial landscapes where recall remains lower and more variable. An additional three-seed representation ablation, run under the original protocol and not repeated after the evaluation corrections, ranked ESM-2 650M highest (mean Recall@2% 0.673 ±0.28) among ProtBert, AntiBERTy, AbLang2, ProGen2, and classical encodings. On BindingGYM, the same configuration matches or exceeds ALLM-Ab on two of three datasets using the 100-sample held-out test metric. Structure-mapped Integrated Gradients and KernelSHAP analyses recover CDR-region attribution patterns consistent with contact-derived paratope geometry, though threshold-free (AUROC) analysis shows this qualitative agreement is not matched by strong residue-level discrimination. These results support Tanimoto-kernel GP active learning as a simple, reproducible strategy for label-efficient antibody affinity optimisation with qualitative structural validation.

## 1 Introduction

Antibody engineering campaigns must choose which candidate mutants to test experimentally from large antibody libraries that contain 10^3^–10^5^ variants (Chen et al., 2016). Biophysical assays such as surface plasmon resonance are highly informative but throughput-limited (Srivastava et al., 2026b)

Active learning (AL) is a strategy for allocating limited assay budget efficiently (Settles, 2012): a cheap surrogate model is trained on the variants labelled so far, then its predictions and uncertainty decide which unlabelled variants to test next (Srinivas et al., 2009), repeating over cycles so a fixed label budget covers more of the library’s high-affinity region than random sampling would. A Gaussian process (GP) is a natural surrogate here, providing a model-based uncertainty estimate for every prediction rather than just a point estimate (Williams and Rasmussen, 2006), which AL needs to avoid getting stuck at a local optimum. Three design choices largely determine how well this loop performs: (1) the numerical representation of each antibody variant, (2) the GP kernel, and (3) the acquisition function that converts the GP’s mean and uncertainty into a rule for selecting the next batch (Gessner et al., 2024; Gorantla et al., 2024).

This paper builds and benchmarks such a framework for antibody-antigen affinity prediction. Antibody variants are represented with ESM-2 650M on the concatenated heavy- and light-chain sequences; the GP surrogate is combined with different kernels and acquisition functions and evaluated on eight public AbBiBench assays (Zhao et al., 2025) against zero-shot and active-learning baselines, under a label-blind seed and a fixed held-out evaluation split (Section 3.2). Because a surrogate is only trustworthy if its reasoning is explainable, we also test whether its learned signal lines up with known antibody-antigen contact geometry via Integrated Gradients (Sundararajan et al., 2017) and KernelSHAP (Lundberg and Lee, 2017) mapped onto deposited crystal structures. The framework reaches 57.8% mean Recall@2% (68.3% on the acquisition-eligible pool), matches or exceeds the AbBiBench zero-shot predictor on all eight datasets, matches or exceeds ALLM-Ab’s GPR(AbLang2) on two of three BindingGYM datasets, and recovers CDR-region attribution patterns consistent with deposited complexes.

We adopt the continuous Tanimoto kernel from Gessner et al. (2024) and confirm its effectiveness across eight datasets; we do not claim this as a new algorithmic contribution. Beyond reproducing this finding, the benchmark reports four empirical observations: (i) linear additivity of the fitness landscape (lin-*R*^2^) predicts AL Recall@2% across libraries at Spearman *ρ* = 0.905 (Section 4.4); (ii) in the original-protocol, three-seed representation ablation, which was not rerun after the evaluation corrections, a general-purpose PLM (ESM-2 650M) ranks above antibody-specific PLMs (AbLang2, AntiBERTy) (Section 4.3); (iii) UCB exploration-schedule choice is essentially irrelevant here, with six schedules spanning only 0.3–0.7 percentage points, replicating an ANOVA finding from small-molecule binding (Srivastava et al., 2026b) (Section 4.5); and (iv) GP predictive-uncertainty calibration flips sign between smooth and rugged fitness landscapes (Section 4.5).

## 2 Related Work

Active-learning protocol choice has been benchmarked in small-molecule affinity settings, where dataset identity often explains more variance than the acquisition rule itself (Gorantla et al., 2024). Srivastava et al. (2026b) extend this direction with an explainable AL framework for ligand binding affinity prediction using GP regression with ECFP/MACCS/ChemBERTa representations and SHAP analysis across four drug targets. They evaluate seven UCB acquisition variants, finding no statistically significant difference between UCB protocols 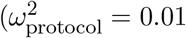, four-factor ANOVA) while all outperform random selection. Dataset identity 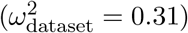 is the dominant factor. Recent representation-learning work also explores assay-supervised graph encoders: GRASP combines molecular-structure pretraining with adaptation on sparse bioactivity assays for molecular-property prediction (Srivastava et al., 2026a); this complements, but is distinct from, our protein-sequence embedding benchmark. A matched-budget study of ionizable-lipid search reports that tested LLM proposers did not significantly outperform genetic-algorithm or random-search baselines (Singh et al., 2026b), further motivating controlled comparisons under fixed assay budgets. Related generative approaches – combinatorial Bayesian optimisation over CDR-H3 sequence space (Khan et al., 2022) and language-model-guided or diffusion-based sequence design (Hie et al., 2021; Stanton et al., 2022; Gruver et al., 2023) – target an open design space, complementary to our pool-based setting over a fixed candidate library.

The eight analysed datasets are drawn from AbBiBench (Zhao et al., 2025), which curates 184,500+ experimental affinity measurements across antibodies and antigens and maintains a public leaderboard of *zero-shot* scoring results. We use their published per-dataset zero-shot Spearman correlations as a citable baseline. Recent work has continued to expand antibody-affinity benchmarking and protein-language-model optimisation protocols, motivating our emphasis on matched datasets, explicit labelling budgets, and reproducible comparisons (Singh et al., 2026a,c). ALLM-Ab (Furui and Ohue, 2025) runs iterative AL for antibody optimisation using GP regression over AbLang2 embeddings (GPR(AbLang2)) - the same model class as ours. We adapt their fixed-budget evaluation protocol (100 seed labels, 12 acquisition cycles of 50 labels, 100-sample held-out test Spearman, 3-seed averaging) on their exact BindingGYM datasets. Gessner et al. (2024) show that a continuous Tanimoto kernel decisively beats stationary kernels (RBF, Matérn) for GP-based antibody affinity AL on PLM embeddings; the Tanimoto/Jaccard similarity itself has a longer lineage as a graph and fingerprint kernel in chemoinformatics (Ralaivola et al., 2005). We adopt their exact kernel formula and confirm the finding on eight diverse analysed datasets. The four validation complexes used for explainability have established paratope contacts in the literature: CR6261/H1N1 HA stem (3GBN) (Ekiert et al., 2009), CR9114/H5N1 HA (4FQI) (Dreyfus et al., 2012; Beukenhorst et al., 2022), Fab-2D1/H1N1 HA head (2FJG) (Xu et al., 2010), and D44.1/lysozyme (1MLC) (Braden et al., 1994). The 3GBN and 4FQI antigen systems are directly represented in our training data enabling in-distribution structural validation.

## 3 Methods

### 3.1 Datasets

The eight analysed assays are taken from the AbBiBench study of experimentally measured antibody-antigen binding data. Each dataset is a combinatorial antibody mutant library screened against a *fixed* antigen (Table 1), covering influenza hemagglutinin stem-binding antibodies CR6261 and CR9114 and AAYL-series scFv libraries against the SARS-CoV-2 Spike HR2 peptide, giving a mixture of small, medium, and large candidate pools.

**Table 1:**
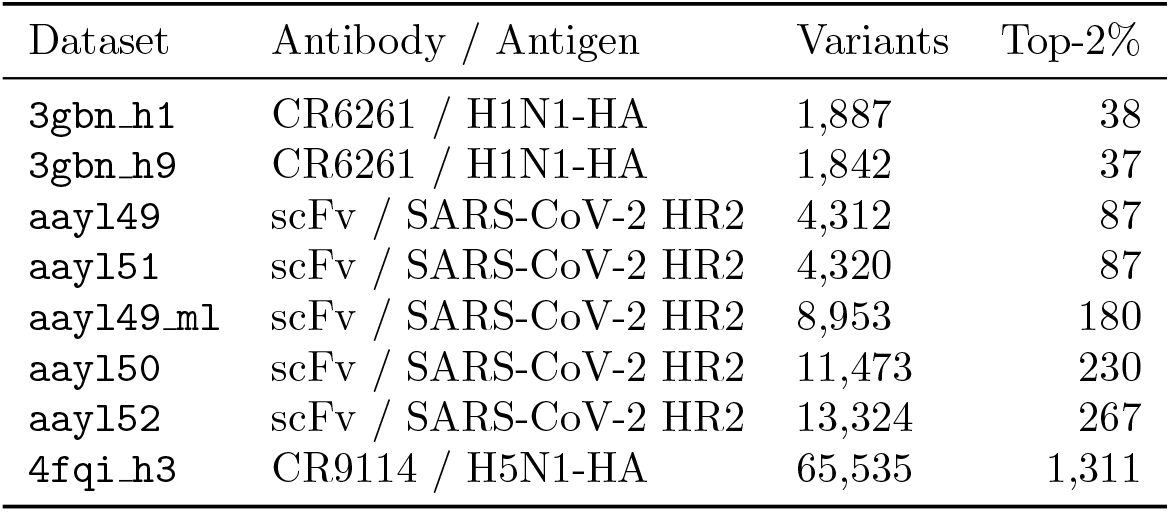
Dataset summary (AbBiBench). HC/LC: heavy/light-chain mutation library. Top-2% counts: top 37-1311 variants per pool.

### 3.2 Active Learning Framework

As demonstrated in Figure 1, we use a pool-based batch AL loop throughout (Settles, 2012). The seed set is chosen *label-blind* : KMeans (*k* = seed size) on the unsupervised PCA embedding, one variant per cluster, giving diverse initial exposure without using affinity labels – replacing an earlier affinity-quantile-stratified version that used information unavailable at seed time in a real campaign. A fixed 15% test split is drawn once per dataset before seeding, excluded from seeding and acquisition throughout, and used for all Spearman evaluation, avoiding the bias of scoring on a shrinking remainder pool that acquisition itself depletes of high-affinity variants. Each cycle fits the GP on labelled data, scores remaining unlabelled (non-test) variants by acquisition, and labels the top batch, repeating until budget exhaustion. We report Recall@2% both against the full-pool top-2% count and against only the top-2% variants eligible for acquisition once the held-out split is removed.

**Figure 1:**
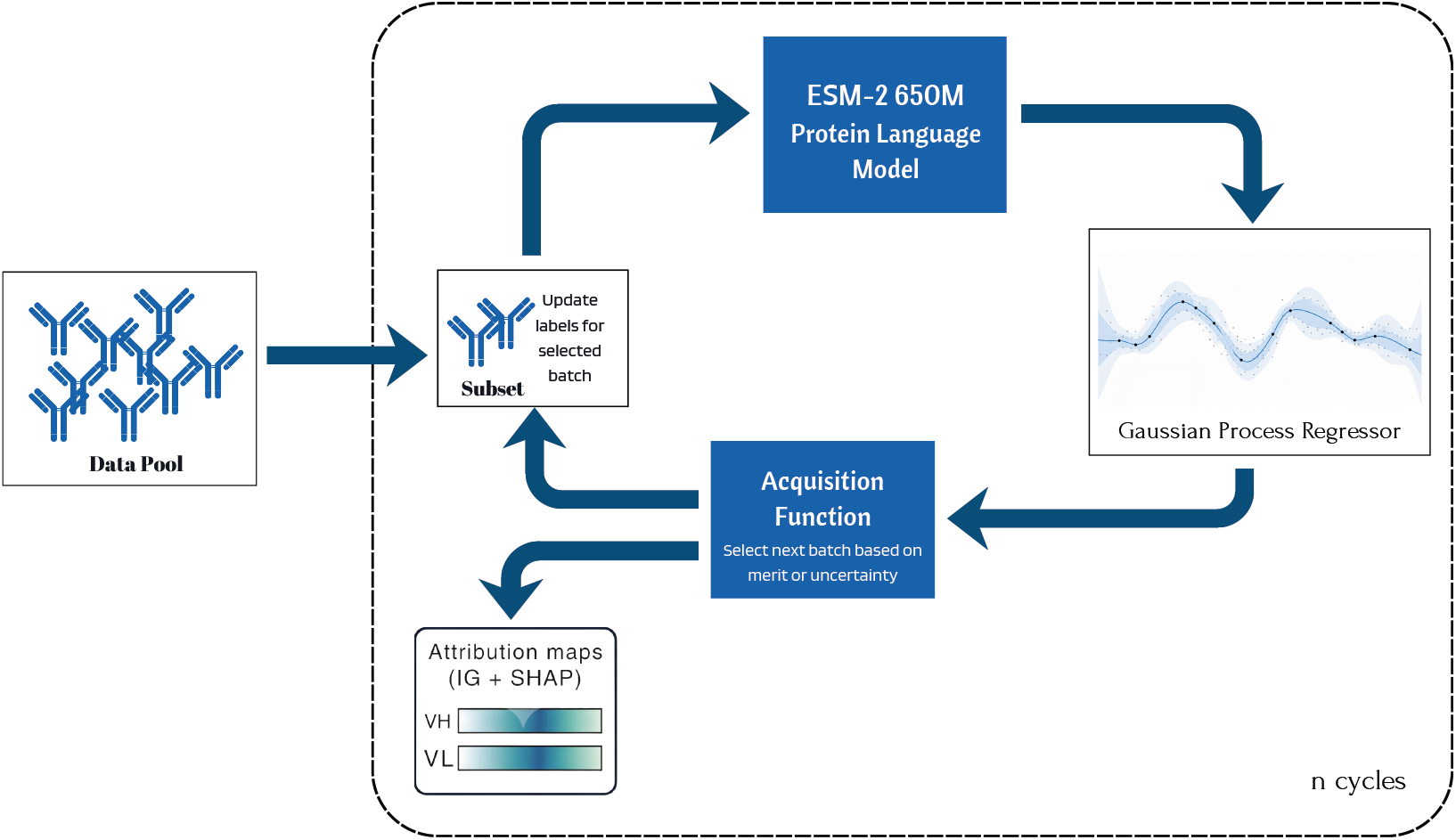
Active-learning workflow. Variants from the data pool are embedded with ESM-2 650M and modelled by a Gaussian process regressor. An acquisition function selects the next batch by merit or uncertainty; labels are updated and returned to the pool for the next cycle. IG and KernelSHAP attribution maps over VH/VL are computed for the best configuration.

### 3.3 Sequence Representation

For each antibody variant, the variable heavy (VH) and variable light (VL) chain sequences are joined with a glycine linker (G) i.e., (VH ∥ `G` ∥ VL) and passed through **ESM-2 650M** (Lin et al., 2023). Per-residue hidden states from the final layer are **mean-pooled** over all positions, yielding a 1,280-dimensional antibody embedding, which is reduced to **128 dimensions** via PCA and standardised.

### 3.4 Gaussian Process Regressor

We use an exact GP (GPyTorch (Gardner et al., 2018; Williams and Rasmussen, 2006)) with a constant mean and compare four kernels. The primary kernel is the continuous Tanimoto kernel,

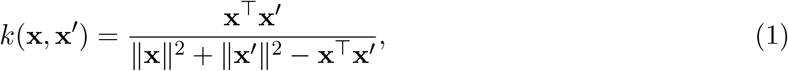

which is scale-invariant, and non-stationary (Gessner et al., 2024). We benchmark it against three standard stationary alternatives: Matérn-5/2,

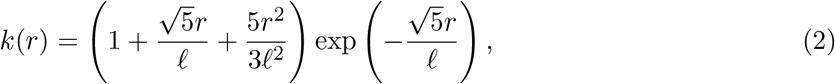

RBF,

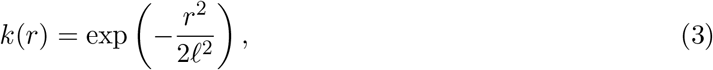

and rational quadratic,

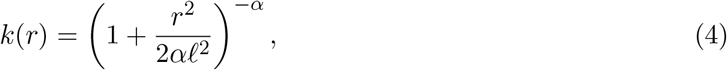

which can be interpreted as a scale mixture of RBF kernels. Hyperparameters are optimised per AL cycle by maximising the exact marginal log-likelihood via Adam (Kingma and Ba, 2017) (100 epochs, lr = 0.1, *γ* = 0.95).

### 3.5 Acquisition Functions

Random selection samples uniformly from the unlabelled pool and serves as the baseline. Following standard Bayesian-optimisation acquistion rules (Srinivas et al., 2009; Jones et al., 1998) Upper confidence bound (UCB) scores each candidate as

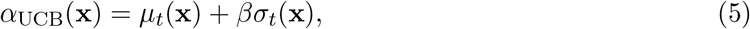

making the exploration-exploitation trade-off explicit through *β*. Expected improvement (EI) uses

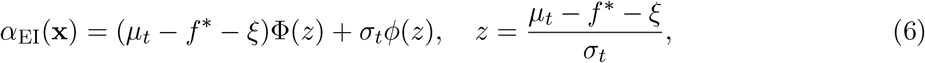

where *f* ^*^ is the best observed value and *ξ* = 0.01. Probability of improvement (PI) uses

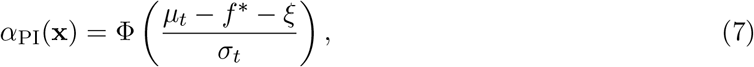

and is therefore the most exploitation-heavy of the non-random strategies. We test six UCB exploration schedules. Fixed schedules use *β* ∈ {0.1, 0.5, 1.0, 2.0}, ranging from exploit-heavy to explore-heavy behaviour. We also evaluate a gradual annealing schedule that decreases linearly from *β* = 2.0 to 0.1, and an alternating schedule that cycles between *β* = 2.0 and 0.1 to explicitly switch between exploration and exploitation. UCB-1.0 (*β* = 1) is used as the standard setting in all other sections.

### 3.6 Labelling Budgets

We adopt a unified 10% label budget across all eight datasets, computed relative to the full pool size *N* before removing the held-out test split. Since 85% of the pool is acquisition-eligible, this corresponds to approximately 11.8% effective coverage of that pool; all reported runs use this full-pool basis, not a 10%-of-eligible budget. For a dataset with *N* variants, the initial labelled set contains *s* = ⌊0.01*N* ⌋ label-blind seed examples. We then run *C* = 9 active-learning cycles, each acquiring *b* = ⌊0.01*N* ⌋ additional measurements, for a final labelled set of *s* + 9*b* ≈ 0.10*N*. This design keeps experimental cost proportional to dataset size and prevents large libraries from dominating the evaluation; Section 4.5 additionally reports a genuinely fixed absolute-label-count comparison across datasets. Unless stated otherwise, all GP hyperparameters are fixed across experiments: PCA dimensionality 128 (fitted on the full pool’s embeddings, unsupervised and label-free, then applied to all variants including the seed set), Adam learning rate 0.1, exponential decay 0.95, 100 epochs per cycle, and UCB *β* = 1.0. Each configuration is repeated with five independent random seeds (42, 123, 456, 789, 1011), and results are reported as mean ± standard deviation. The primary metric is Recall@2%, the fraction of true top-2% binders recovered in the labelled set; we also report Spearman *ρ* between the GP predictive mean and true affinity on the held-out test split. GP training and its marginal-likelihood objective are exact; predictive standard deviations (used by UCB/EI/PI and by the calibration analysis in Section 4.5) use GPyTorch’s `fast_pred_var` approximation, and datasets whose labelled set exceeds GPyTorch’s default `max_cholesky_size` of 800 (`aayl49_ml`, `aayl50`, `aayl52`, `4fqi_h3`, on later cycles) use an iterative conjugate-gradient solver rather than direct Cholesky factorisation for both fitting and prediction.

### 3.7 Explainability: IG and KernelSHAP

We compute per-residue attributions for the best configuration (Tanimoto + UCB) using two complementary methods.

#### Integrated Gradients (IG)

(Sundararajan et al., 2017) **integrates the gradient of the GP predictive mean *µ***_***t***_ **along a straight path from an all-**`<mask>` **baseline x**′ to the input **x**:

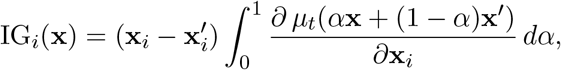

approximated by a 50-step Riemann sum. Attribution for residue *i* is the *l*_2_ norm of the 1,280-dimensional gradient vector at that position, collapsed to a scalar.

#### KernelSHAP

(Lundberg and Lee, 2017) treats each residue position as a binary feature (present or masked), estimating Shapley values via kernel regression with 256 background samples. **Ground-truth contacts** are computed from deposited PDB coordinates for four systems with independently published crystal structures and literature-established paratopes (3GBN (Ekiert et al., 2009), 4FQI (Dreyfus et al., 2012; Beukenhorst et al., 2022), 1MLC (Braden et al., 1994), 2FJG (Xu et al., 2010)), under five definitions: (i) heavy-atom distance ≤ 4.5 Å (Biopython NeighborSearch), (ii) a ±1-residue-tolerant version of (i), (iii) the literature-cited paratope, (iv) a ±1-residue-tolerant version of (iii), and (v) an extended 8 Å contact. We report AUROC under all five, with attributions mapped to structure residue numbers via BLOSUM62 alignment. IG and KernelSHAP require many repeated differentiable passes per residue per variant, tractable only on a small probe set; we fit this probe GP on a random 800-point subsample per dataset and verify it is a strong predictor (held-out Spearman *ρ* = 0.974 on `3gbn_h1`) before attributing its decisions. The 3GBN/4FQI probe GPs are in-distribution (trained on the same antigen system); 1MLC/2FJG are out-of-distribution controls. A separate six-complex analysis additionally includes four AAYL model structures provided by AbBiBench, without independent literature paratope citations.

## 4 Results

### 4.1 Tanimoto+UCB is the Best Point-Estimate Active-Learning Configuration

Figure 2 summarises performance across the eight datasets under the unified 10% budget, label-blind seeding, and held-out evaluation (Section 3.2). Across all 16 kernel-protocol configurations, Tanimoto+UCB achieves the highest mean Recall@2% (0.578, or 0.683 against the acquisition-eligible pool), followed by RQ+UCB (0.567), Tanimoto+EI (0.557), and RQ+EI (0.554), all substantially above random selection (0.096). This AL-versus-Random gap is significant by a paired Wilcoxon signed-rank test across datasets (*p* = 0.0078, the floor at *n* = 8, using seed-averaged Recall@2% per dataset as the unit of analysis). The finer ranking among active configurations is not distinguishable at this sample size (Tanimoto+UCB vs. RQ+UCB, *p* = 0.44; vs. Tanimoto+EI, *p* = 0.09), so we report Tanimoto+UCB as the best point estimate, not a significantly superior configuration.

**Figure 2:**
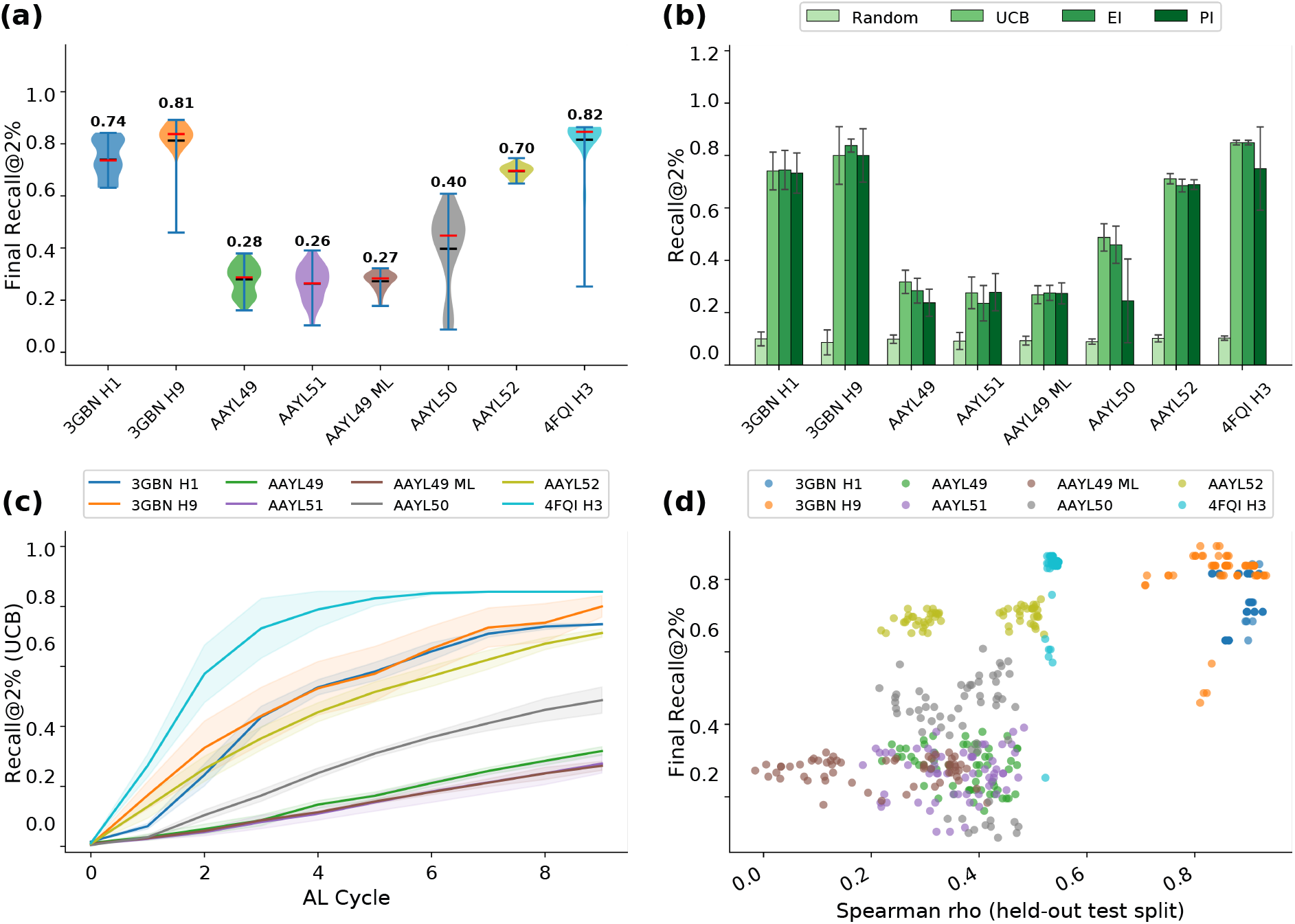
Overview of active-learning performance across eight antibody-antigen targets (ESM-2 650M, VH+G+VL concatenation, unified 10% label budget, five seeds). (a) Final Recall@2% distribution across all 16 kernel-acquisition combinations. (b) Per-target Recall@2% by acquisition protocol (mean ± std, 4 kernels, five seeds). (c) UCB learning curves (mean ± std, 4 kernels, five seeds). (d) Recall@2% vs. Spearman *ρ* across all kernel-protocol-dataset combinations.

Figure 3 expands the overview by target, showing protocol-kernel bars and active-learning trajectories for UCB, EI, and Random. Per-dataset results reveal two regimes. Structurally constrained single-chain influenza libraries exhibit the largest recovery: Tanimoto+UCB recovers 74.2%, 82.7%, and 84.9% of top-2% binders on `3gbn_h1`, `3gbn_h9`, and `4fqi_h3`, respectively (89.2%, 98.1%, and 100.0% against the acquisition-eligible pool). The AAYL combinatorial libraries are harder: `aayl49` reaches 33.6%, `aayl49_ml` 27.3%, and `aayl51` 31.0%, while `aayl50` (56.0%) and `aayl52` (72.7%) recover substantially more despite comparable combinatorial complexity (Section 4.4 relates this spread to landscape epistasis).

**Figure 3:**
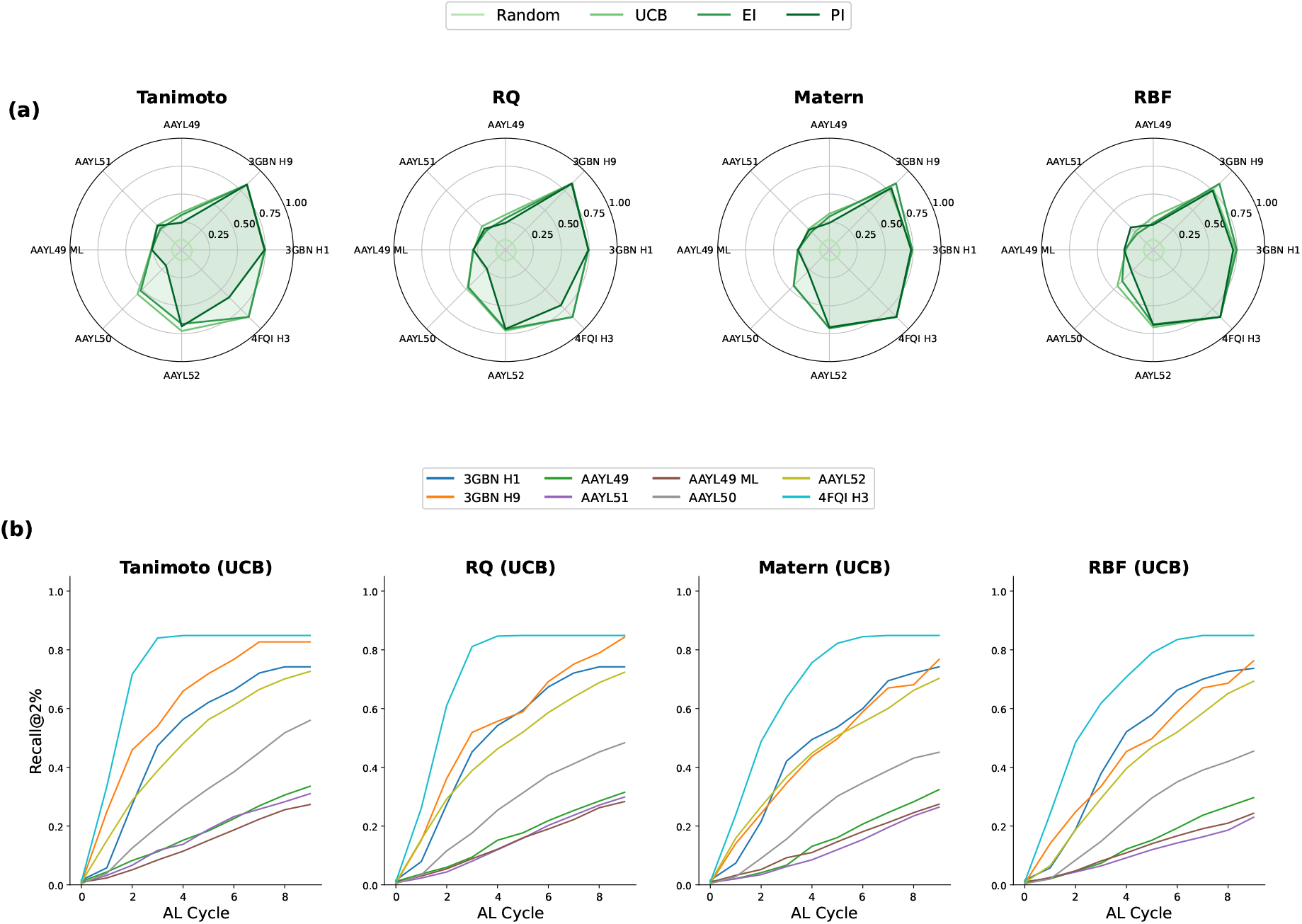
Per-target ablation views across all 8 antibody-antigen datasets under the unified 10% budget, faceted by kernel. (a) Radar plots of final-cycle Recall@2% across all 8 datasets (axes), one polygon per acquisition protocol, one subplot per kernel. (b) Active-learning curves (Recall@2% vs. AL cycle; mean across five seeds, UCB acquisition), one line per dataset, one subplot per kernel.

The compact kernel summary in Figure 5a confirms the same design choice from a different aggregation: Tanimoto is the best point estimate on UCB and EI, with UCB reaching 0.578 mean Recall@2%, EI 0.557, PI 0.484, and Random 0.096. Among stationary kernels, RQ is consistently competitive with Tanimoto and outperforms Matérn and RBF, though none of these differences are statistically significant at 5 seeds (Section 4.1). The UCB-Random gap (0.482) exceeds the Tanimoto-RBF gap within UCB (0.045), indicating that acquisition strategy matters more than kernel choice when deploying GP active learning for antibody optimisation.

### 4.2 Comparison Against Prior Work

Table 2 compares Spearman *ρ* on the fixed held-out test split (Section 3.2), never seen during training or acquisition, correcting an earlier full-pool evaluation. We additionally report GP+Random at the same budget to separate the effect of having labels at all from active acquisition specifically. GP+UCB achieves a mean *ρ* of 0.548 versus 0.435 for the best zero-shot baselines, matching or exceeding it on all eight datasets (within seed noise on `aayl49` and `aayl49_ml`). GP+Random is at least as good as GP+UCB on seven of eight datasets, as expected: UCB’s exploitation bias helps Recall@2% but not full-range rank correlation. The improvement over zero-shot thus comes mainly from having any labels at all; UCB’s specific advantage is top-binder discovery under a cost-matched budget (Section 4.5).

**Table 2:**
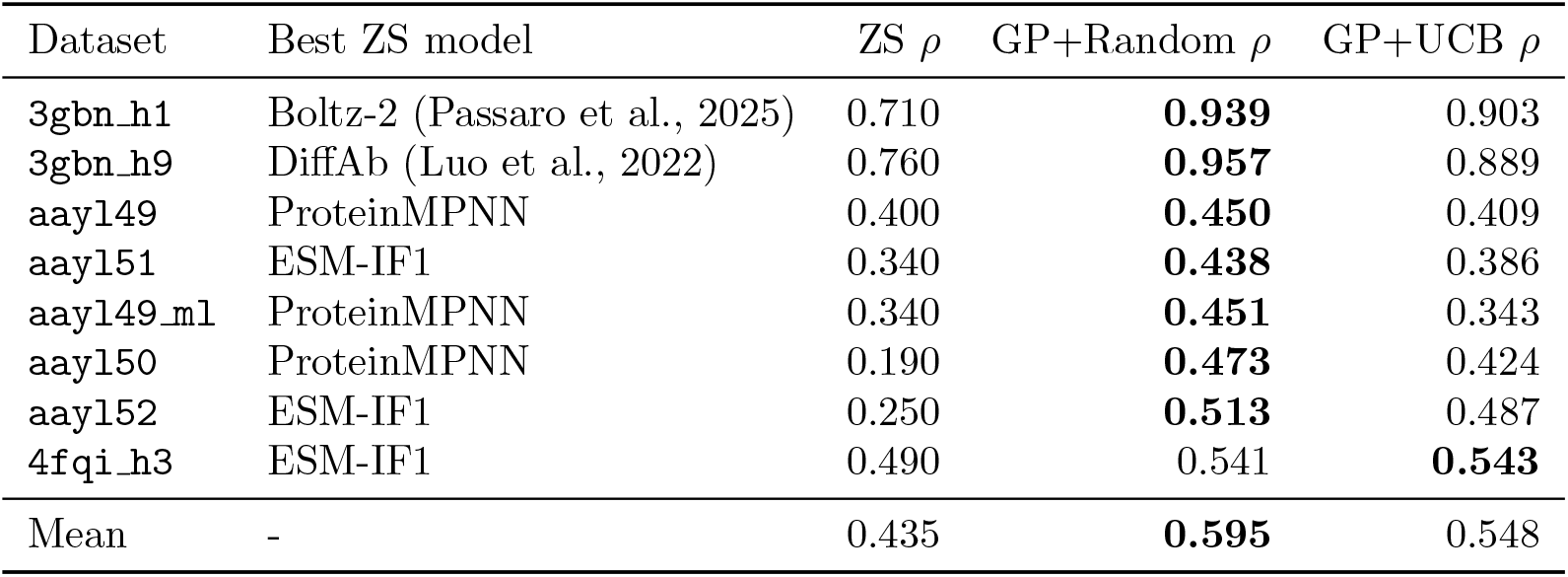
AbBiBench best zero-shot (ZS) Spearman *ρ* vs. our framework, evaluated on the held-out test split. ZS scores are the best reported values from AbBiBench (Zhao et al., 2025); Protein-MPNN (Dauparas et al., 2022) and ESM-IF1 (Hsu et al., 2022) are the inverse-folding ZS baselines appearing below. GP columns use ESM-2 650M + Tanimoto kernel with ≈10% of each pool labelled over nine active-learning cycles (mean, five seeds). ↑ indicates higher is better; bold marks the row maximum. GP+Random matches or exceeds GP+UCB on this full-range ranking metric on seven of eight datasets (Section 4.2), while both GP variants match or exceed the best ZS model on all eight.

ALLM-Ab (Furui and Ohue, 2025) benchmarks GPR(AbLang2) on BindingGYM datasets and identifies it as the strongest active-learning surrogate among their compared methods. We adapt their fixed-budget protocol: 100 seed labels followed by 12 acquisition cycles of 50 labels (700 labels total), Spearman *ρ* evaluated on a 100-sample held-out test set drawn and excluded from the pool before seeding (this pipeline uses plain uniform random seed sampling throughout and never carried the affinity-stratified seeding issue described in Section 3.2), and three-seed averaging. Under this matched test-set metric, Tanimoto+UCB matches or exceeds GPR(AbLang2) on two of three datasets without fine-tuning: on `5A12_Ang2`, our method reaches *ρ* = 0.537 ± 0.100 versus GPR(AbLang2) at ≈ 0.33 and ESM2 FT at ≈ 0.25; on `5A12_VEGF`, our method reaches 0.655 ± 0.054 versus ≈ 0.60 for GPR(AbLang2) and ≈ 0.47 for ESM2 Finetune. On `4D5_HER2`, our result (0.422 ± 0.191) is numerically below both baselines (≈ 0.45), with a standard deviation wider than the gap – comparable, not exceeding. This BindingGYM comparison uses antibody-affinity data from a separately published, independently deposited source, distinct from AbBiBench, and is therefore also our clearest held-out generalisation evidence (Section 5).

### 4.3 ESM-2 Leads the Original-Protocol Representation Ablation

We evaluate representation choice in two complementary ways: first by comparing six protein language models under the same GP pipeline, and then by replacing PLM embeddings with classical sequence featurisations. These ablations use the original three-seed protocol and were not rerun under the corrected seeding and held-out evaluation. Figure 4a shows that ESM-2 (Lin et al., 2023) ranks highest in this ablation (0.673 ± 0.28 mean Recall@2%), followed by ProtBert (Elnaggar et al., 2021) (0.627 ± 0.31), paired AntiBERTy (Ruffolo et al., 2021) (0.615 ± 0.28), AbLang2 (Olsen et al., 2024)(0.604 ± 0.31), concatenated AntiBERTy (0.579 ± 0.32), and ProGen2 (Nijkamp et al., 2023)(0.507 ± 0.31). Figure 4b shows that classical features remain weaker: bag-of-amino-acid (Leslie et al., 2001) 5-mers reach 0.581 ± 0.32, BLOSUM62 (Henikoff and Henikoff, 1992) reaches 0.520 ± 0.32, and one-hot encoding reaches 0.524 ± 0.31.

**Figure 4:**
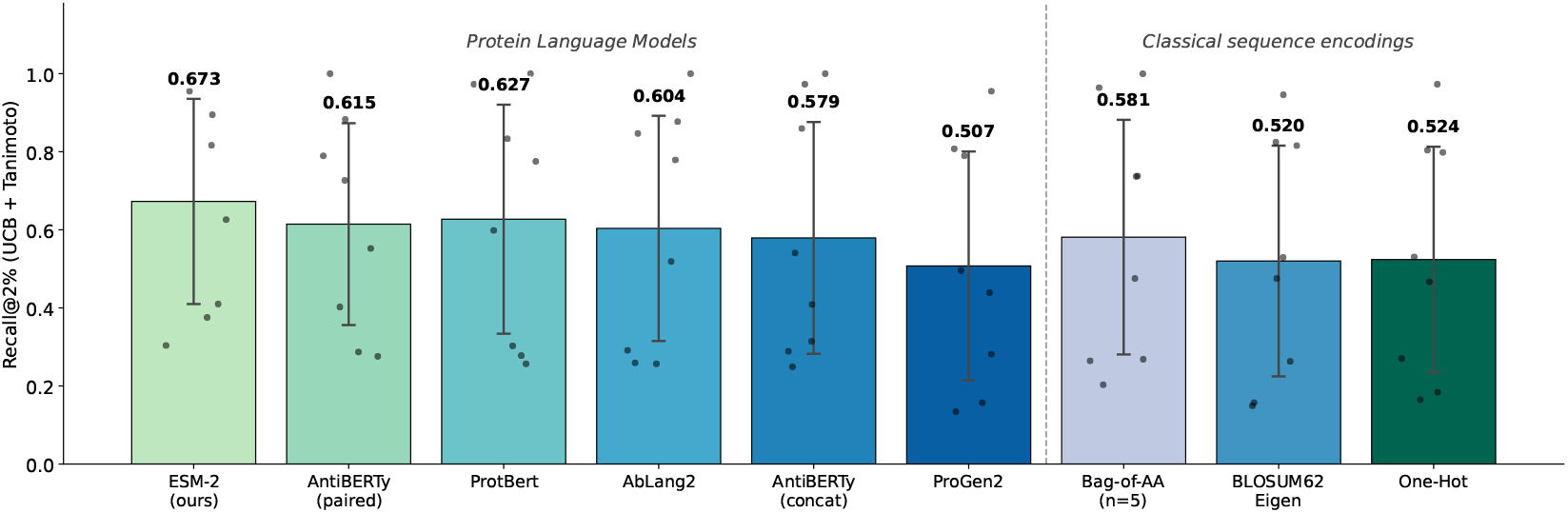
Representation ablation: mean Recall@2% ± std across 8 targets (Tanimoto + UCB, unified 10% budget, original protocol, three seeds (not rerun under the corrected protocol); dots = individual targets; error bars = cross-target std). (a) PLM comparison. ESM-2 650M achieves the highest mean (0.673 ± 0.28) despite not being antibody-specific, followed by ProtBert (0.627 ± 0.31), AntiBERTy-paired (0.615 ± 0.28), AbLang2 (0.604 ± 0.31), AntiBERTy-concat (0.579 ± 0.32), and ProGen2 (0.507 ± 0.31). (b) Featurisation comparison. Bag-of-amino-acid 5-mers are the strongest classical baseline (0.581 ± 0.32), but remain 9.2 pp below ESM-2. One-hot encoding (0.524 ± 0.31) and BLOSUM62 eigendecomposition (0.520 ± 0.32) perform similarly and lack the contextual, co-evolutionary signal captured by large PLMs.

**Figure 5:**
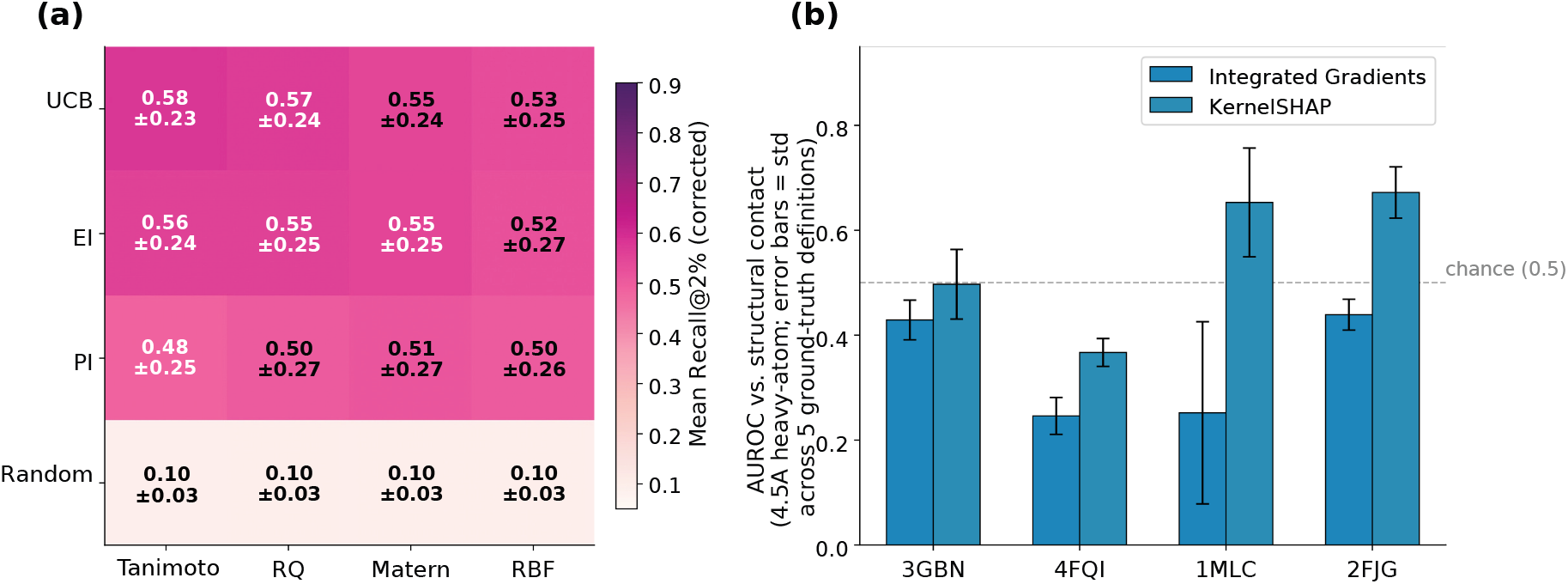
Kernel selection and structural explainability summary, corrected protocol. (a) Kernel × acquisition heatmap (mean ± std Recall@2%, 8 targets, ESM-2 concat, unified 10% budget, five seeds). Tanimoto is the best point estimate on UCB (0.58) and EI (0.56), followed closely by RQ, with Matérn and RBF close behind and Random (0.10) far below; differences among active configurations are not statistically significant (Section 4.1). (b) AUROC of attribution score against the 4.5 Å structural-contact label (threshold-free, continuous ranking) for Integrated Gradients (IG) and KernelSHAP across the four systems with independently published crystal structures. IG’s AUROC is at or below chance (0.5) on all four systems; KernelSHAP is above chance on 1MLC and 2FJG, where VL contributions are substantial, and near chance on 3GBN/4FQI.

ESM-2 650M outperforms all alternatives despite not being antibody-specific, plausibly because its scale and broad pre-training corpus produce an embedding geometry that aligns well with the Tanimoto kernel’s inner-product similarity measure; ProGen2’s weaker performance may reflect its autoregressive objective being less suited to discriminative GP regression. Among classical encodings, bag-of-AA pentapeptide k-mers are strongest, but the persistent 9.2 pp gap to ESM-2 indicates contextual PLM features matter for non-additive CDR landscapes.

### 4.4 Influenza Libraries Saturate Early but AAYL Landscapes Need Larger Budgets

Budget-curve analyses show Recall@2% vs. % of dataset labelled, using UCB acquisition averaged across four kernels. Influenza datasets (`3gbn_h1`, `3gbn_h9`, `4fqi_h3`) reach their strongest recall (74.2–84.9%) by the full 10% budget. AAYL datasets improve more gradually: `aayl50`/`aayl52` reach 56.0%/72.7%, while `aayl49`, `aayl49_ml`, and `aayl51` plateau below 35%, indicating that simultaneous VH+VL combinatorial libraries are fundamentally harder for GP regression under a fixed budget (Weinreich et al., 2006; Poelwijk et al., 2019). We quantify this via nearest-neighbour fitness correlation (*r*_nn_) and linear additivity (lin-*R*^2^; 5-fold ridge regression on the ESM-2 embeddings), where low lin-*R*^2^ indicates epistasis (Poelwijk et al., 2019). Influenza libraries are near-additive (lin-*R*^2^ = 0.785, *r*_nn_ = 0.844) versus rugged AAYL libraries (lin-*R*^2^ = 0.228, *r*_nn_ = 0.356); lin-*R*^2^ predicts Recall@2% at Spearman *ρ* = 0.905 across all eight datasets, a practical pre-screen for landscape amenability before committing to a full AL campaign.

### 4.5 Computational Cost, Fixed Budgets, Calibration, and Acquisition Scheduling

#### Computational cost

GP training scales 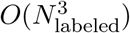 per cycle, not 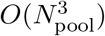. On a single RTX A4000, full 10-cycle AL runs took 7–76 s depending on pool size (largest: `4fqi_h3`, 6,550 labelled points at the final cycle), with peak GPU memory 3.0 GB, well under 1 GB for the small datasets. The 800-point probe GP used for explainability is a separate, smaller model and is not part of this cost.

#### Fixed absolute budget

The unified 10% budget spans approximately 190–6,550 labels, so we also ran an equal-cost schedule (100 seed + 10×50 = 600 labels per dataset, Tanimoto kernel, three seeds), evaluated against the maximum recall attainable at 600 labels (full table in the supplementary information). GP+UCB exceeds GP+Random on every dataset (paired Wilcoxon *p* = 0.0078; e.g., `3gbn_h9`: 84.7% vs. 37.8%). For `4fqi_h3`, the ceiling is 38.3%; its 34.4% recall is 89.8% of that maximum. The other seven datasets have a 100% ceiling.

#### Uncertainty calibration

We computed 90% credible-interval coverage on the held-out test split at the final cycle (*n*_test_ per dataset is exactly 15% of its pool). The GP is *overconfident* on the near-additive influenza libraries (`3gbn_h1`: 82.7%, `3gbn_h9`: 81.5%, against the 90% nominal target) and *conservative* on the epistatic AAYL libraries (`aayl49`: 98.5%, `aayl51`: 95.8%, `aayl49_ml`: 96.2%, `aayl50`: 98.1%, `aayl52`: 98.5%, `4fqi_h3`: 96.5%). Restricted to the five AAYL datasets, mean Recall@2% spans only 43.3–44.0% across all six *β* schedules below, so an exploit-heavy schedule that discounts uncertainty does not recover more recall – suggesting the GP’s predictive mean itself carries little usable ranking signal on these landscapes, independent of how its uncertainty is weighted. A per-mutation residual analysis proposed during rebuttal was not completed for this version and remains future work.

#### UCB *β*-scheduling

Section 3.5 describes six exploration schedules; we report the complete ablation here (6 schedules × 8 datasets × 3 seeds): fixed *β* ∈ {0.1, 0.5, 1.0, 2.0} reach 57.5%, 57.2%,57.4%, and 57.4% mean Recall@2% (44.0%, 43.5%, 44.0%, 43.7% on the 5 AAYL datasets), while the gradual (2.0→0.1) and alternating (2.0/0.1) schedules reach 57.3% and 57.2% (43.7%, 43.3% on AAYL). All six fall within 0.3–0.7 percentage points of one another, both overall and on the harder AAYL subset: *β* choice does not matter meaningfully in this setting. This independently replicates, in antibody affinity active learning, the finding of no significant UCB-protocol effect reported for small-molecule binding by Srivastava et al. (2026b).

### 4.6 Attributions Emphasise CDR Regions but Lack Residue-Level Discrimination

We use four crystal structures to assess whether IG and SHAP attributions are biologically plausible when mapped back onto antibody-antigen complexes and compared with known paratope and epitope geometry.

The two influenza stem systems, CR6261/H1N1 HA (3GBN) and CR9114/H5N1 HA (4FQI), are represented in the AbBiBench training data and therefore test in-distribution structural consistency. D44.1/lysozyme (1MLC) and Fab-2D1/H1N1 HA head (2FJG) provide out-of-distribution and cross-epitope controls.

Across these cases, the attributions consistently emphasise CDR-region structure and chain participation patterns compatible with established structural ground truth (Ekiert et al., 2009; Dreyfus et al., 2012; Xu et al., 2010; Braden et al., 1994; Beukenhorst et al., 2022): the 3GBN and 4FQI interfaces concentrate contacts around compact antibody-HA stem regions (CR6261/3GBN VH-dominated; CR9114/4FQI strongest around the HCDR2-centred stem interface). On the four systems with independently published crystal structures (Figure 6), IG’s AUROC against the structural-contact label is at or below chance on all four (0.43, 0.25, 0.25, 0.44 for 3GBN/4FQI/1MLC/2FJG), holding under the extended contact definition and both tolerance versions. KernelSHAP is more mixed: near chance on 3GBN/4FQI (0.50, 0.39) but above chance on 1MLC/2FJG (0.65, 0.67), where substantial VL contributions let SHAP’s feature-masking better capture inter-chain dependencies that IG’s gradient diffuses across the mean-pool operation.

**Figure 6:**
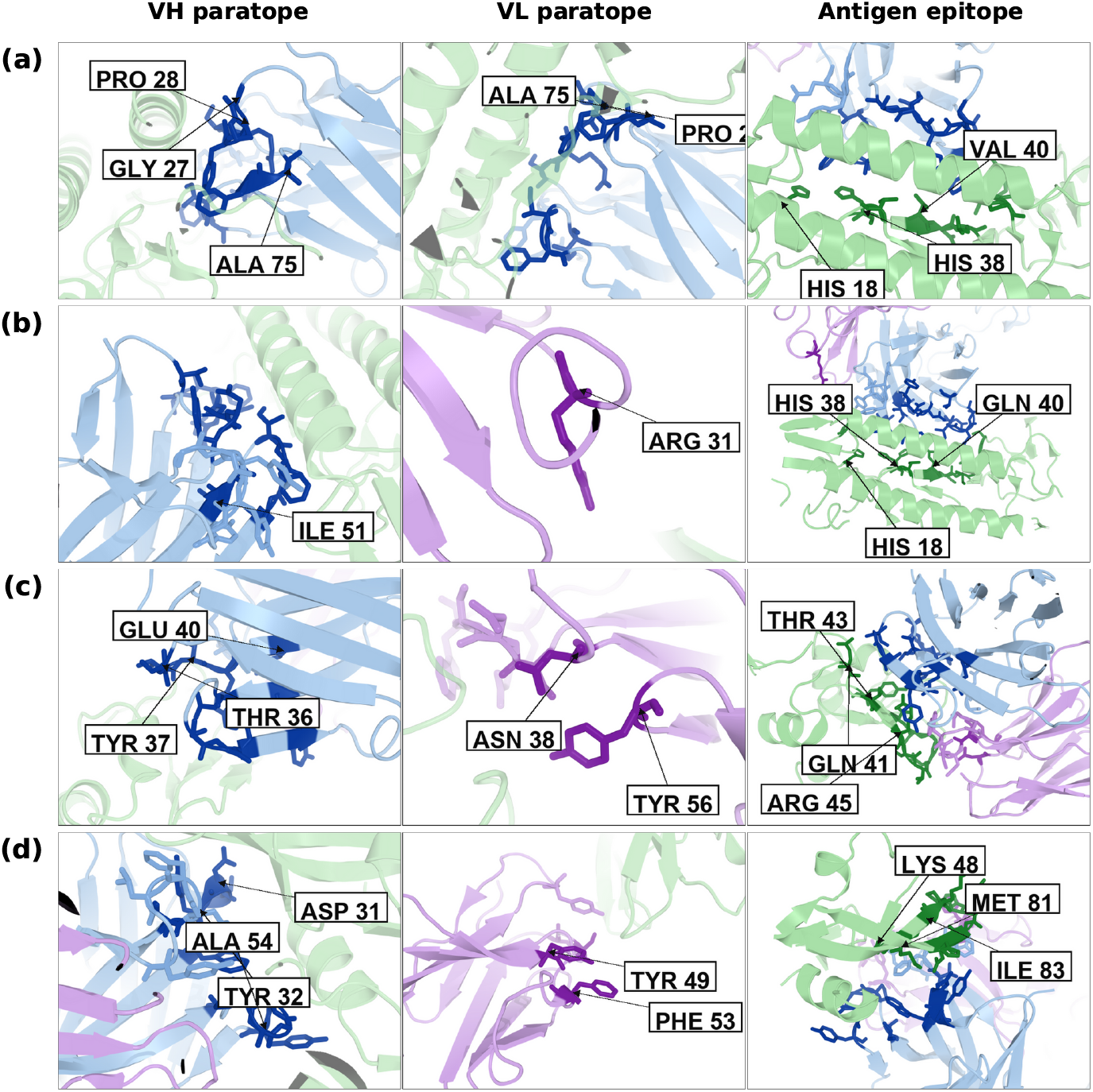
Structural validation and ground-truth contact panels for the explanation case studies. Contact-derived paratope and epitope residues computed from deposited coordinates provide the reference used to evaluate whether attribution maps align qualitatively with known binding modes.(a) CR6261/H1N1 HA stem (3GBN) shows VH-dominated stem recognition, with attribution concentrated near heavy-chain CDR regions and little VL signal, consistent with the known binding mode. (b) CR9114/H5N1 HA (4FQI) highlights the HCDR2 region and residues around the known influenza stem interface, providing the clearest in-distribution qualitative agreement. (c) D44.1/lysozyme (1MLC), an out-of-distribution control, still shows CDR-centred heavy- and light-chain attribution patterns compatible with lysozyme recognition. (d) Fab-2D1/H1N1 HA head (2FJG), a cross-epitope control, shows dual-chain attribution consistent with head-site recognition rather than the CR6261 stem mode. Together, these panels support qualitative CDR-region explainability while cautioning against residue-exact claims from mean-pooled embeddings.

A separate six-complex analysis additionally includes AAYL49/50/51/52, four SARS-CoV-2 HR2 scFv systems with AbBiBench-provided model structures rather than independently deposited crystal structures; we distinguish these two structural-evidence tiers explicitly. Attributed residues concentrate in or near CDR loops across all four literature-validated panels, including the out-of-distribution 1MLC/2FJG controls, but the AUROC analysis rules out a stronger, residue-exact reading, since mean-pooling the ESM-2 embedding before the GP discards positional information while still ranking affinity well (Spearman up to 0.90, Table 2) using diffuse, non-localised signal. Together, these results support qualitative, CDR-region-level structural validation, but not residue-level discrimination.

## 5 Discussion and Conclusions

Across eight AbBiBench libraries, the five-seed main grid gives Tanimoto+UCB the highest mean point estimate (57.8% full-pool and 68.3% acquisition-eligible Recall@2%), significantly above Random (*p* = 0.0078), including under a fixed 600-label budget, but not significantly different from RQ+UCB or Tanimoto+EI. The UCB–Random gap exceeds the Tanimoto–RBF gap under UCB. The original-protocol, three-seed representation ablation ranked ESM-2 highest, with 5-mers the strongest non-PLM baseline; this comparison was not rerun after protocol corrections.

On independently sourced BindingGYM, our model matches or exceeds ALLM-Ab’s GPR(AbLang2) on two of three datasets and is indistinguishable on the third under its 100-sample held-out test metric, providing external generalisation evidence.

The BindingGYM CD19/FMC63 dataset (PDB 7URV) varies antigen sequence only; FMC63 is identical across all 3,886 rows. No other BindingGYM assay varies antibody VH/VL, so the three datasets above are the available held-out antibody-mutation evidence. Prospective validation on new antibody families remains future work.

IG and KernelSHAP emphasise CDR-region structure across four crystal-structure case studies, including in-distribution and control systems. However, AUROC does not support residue-level discrimination: IG is at or below chance throughout, while KernelSHAP is mixed. This in-silico study measures retrospective label recovery, not prospective experimental efficiency under assay noise. The datasets cover few antibody-antigen families, and BindingGYM is the only independent generalisation evidence. Structural validation uses four deposited crystal structures and assesses attribution agreement, not causal explanations of predictions.

Overall, we position this work as a benchmark, not a new algorithm, and identify ESM-2, a Tanimoto-kernel GP, UCB, and a 10%-of-full-pool budget as a practical default. Epistatic VH+VL libraries may require larger budgets or higher-capacity surrogates. Future work includes multi-antigen transfer, prospective validation, and generative design; structural attributions should remain qualitative CDR-region guidance.

## Acknowledgements

Harshit Singh would like to acknowledge the support of the Department of Computer Science and Engineering, Shiv Nadar University, Delhi-NCR, India, for providing all the resources needed for computation.

## Data and Code availability

Source code and scripts are available at https://github.com/harshitsinghsnu/AntibodyAL. AbBiBench datasets: https://huggingface.co/datasets/AbBibench/Antibody_Binding_Benchmark_Dataset

BindingGYM datasets: https://zenodo.org/records/12514160.

## A Experiment Configuration

**Software**. GPyTorch 1.15.2 / PyTorch 2.11.0 (CUDA); ESM-2 650M via fair-esm 2.0.0; Biopython 1.83; PyMOL Open-Source 3.1.0. **Hardware**. NVIDIA RTX A4000. **PCA**. 128 components, fitted on the full pool’s ESM-2 embeddings, applied to all variants including the seed set. **GP**. Constant mean; 100 training epochs; Adam, lr = 0.1; *γ* = 0.95; prediction batch size 5,000 with fast pred var (predictive SD only; training is exact). GPyTorch’s default max cholesky size = 800 is not overridden, so labelled sets above 800 points use an iterative conjugate-gradient solver. **Seed selection**. Label-blind KMeans (*k* = seed size) on the unsupervised PCA embedding, one variant per cluster. **Evaluation split**. Fixed 15% test split, drawn before seeding and excluded throughout. **Budget**. 10% of full pool (before holdout): 1% seed + nine 1% cycles; fixed 600-label comparison also reported. **Seeds**. 42, 123, 456, 789, 1011. **Code**. Source code and scripts are available at https://github.com/harshitsinghsnu/AntibodyAL. **Compute**. See Section 4.5. **Total configurations**. 640 main (4 *×* 4 *×* 8 *×* 5) + 144 PLM + 72 representation + 144 *β*-ablation + 48 fixed-budget runs.

